# The swarming behaviors of Vorticella

**DOI:** 10.1101/2024.11.27.625613

**Authors:** Tetsuya Tabata, Yuko Maeyama

## Abstract

Two Vorticella species undergo a synchronous transition from sessile zooids to motile telotrochs, which swarm enveloped in secreted mucus, subsequently forming dense aggregations on substrates and reverting to the zooid form. This cyclical process recurs on a daily basis. Each species exhibits a unique mode of swarming behavior. We hypothesize that these behaviors may serve to facilitate efficient feeding while concurrently acting as a mechanism for predator avoidance.

## Introduction

“It has long been known that animals may aggregate into groups or clusters, more or less closely associated, in which physical contact may, or may not, occur. Normally actual physical contact is found as a part of the aggregation phenomenon in many Protozoa, Paramecium for example; in flatworms such as the planarians; in earthworms; in arthropods and molluscs; and among many vertebrates, including fish, frogs, reptiles, birds and mammals” (Allee, 1927) In this study, we present further examples within the ciliate protozoan species, Vorticella spp., demonstrating an alternation between two distinct life cycle phases: the sessile, stalked-feeding zooid and the motile swarmer, known as the telotroch. This transition is typically attributed to environmental stressors (Barlow and Finley, 1976). However, in this study, we demonstrate that in these particular species, the transition occurs cyclically on a daily basis, wherein zooids synchronously transform into telotrochs, which subsequently swarm and collectively settle or congregate on solid substrates. We had recognized this behavior has only been publicly documented in conference abstracts by the research group of Horikami and Ishii, one of which was in the international meeting where they reported that four Vorticella species exhibit this type of behavior (Horikami and Ishii, 1981). During manuscript preparation, we also uncovered earlier reports describing the gregarious Vorticella species by Fox and Newth (Fox and Newth, 1936; Fox and NEWTH, 1935). Our current study confirms their findings and presents the gregarious behaviors using videos, a method not yet feasible in earlier pioneering research.

## Results and Discussion

### Swarming behaviors of Vorticella

We identified two species of the genus *Vorticella* from Sanshirou-ike, the pond, situated within the University of Tokyo campus. The pond’s name originates from a famous novel by Souseki Natsume, written during the Meiji era. We primarily focus on the first species, the provisionally named *Vorticella sp11* (Sanshirou) (Figure 1) as its ITS1-5.8S-ITS2 DNA sequence is identical to that of *Vorticella sp11* (Figure 1-supplementary figure1), a species originating from China that has been solely characterized by its genetic sequence (Sun et al., 2013). *Vorticella sp11* (Sanshirou) appeared seasonally, from late April to early July. Despite efforts, we were unable to establish stable culture conditions and could only maintain the organisms for a few weeks post-sampling. Periodic sequencing of samples was performed to verify the genotype of the observed individuals. Although not all samples were sequenced, those displaying the characteristic behaviors of V. sp11 (Sanshirou) consistently exhibited identical ITS1-5.8S-ITS2 DNA sequences, without exception, despite the sequence being relatively short at approximately 400 base pairs. We acknowledge that species identification based solely on morphology was not possible, and thus we cannot exclude the possibility that the individuals presented as *V. sp11* (Sanshirou) in this study may encompass heterogenous species.

**Figure 1.**
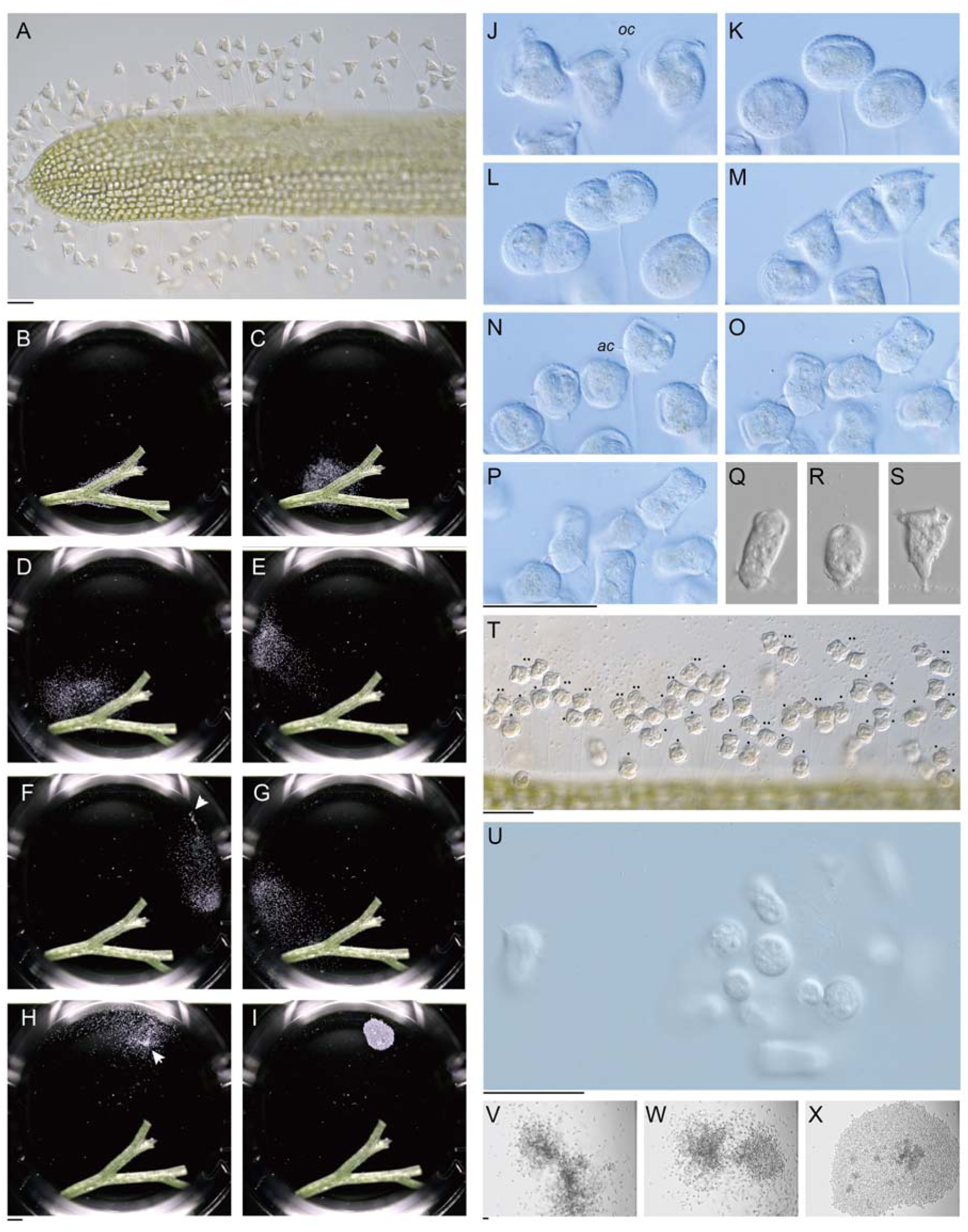
Life cycle of *V. sp11* (Sanshirou). (A) Zooids affixed to a *Cabomba sp.* leaf. (B-I) The transition from zooids (B) to telotrochs (C) and their subsequent group settlement (I). Arrows denote the condensed mucus (F, H). (J-P) Cellular division and the ensuing transition from zooids (J) to telotrochs (O). (J) *oc* denotes oral cilia, while (N) *ac* denotes aboral cilia. (Q-S) Telotroch settlement and subsequent transition to a zooid form. (T) Telotroch formation via cellular division (..) and in its absence (.). The video content is identical to that in (A) but presented in a vertically inverted orientation. (U) The settlement process is shown in high-resolution video recordings. (V-X) An instance of the settlement process involving a substantial number of telotrochs. Scale bars represent 100 μm in all instances, except for (B-I), where the scale bar denotes 1 mm. All figures represent frames from video recordings, with online access to corresponding video images provided below and hereafter. **Figure 1-video 1 (related to Figure1 B-I)** The transition from zooids to telotrochs and their subsequent group settlement. The playback rate is accelerated by a factor of 100 in select segments. https://youtu.be/iCczJqXLKbA **Figure 1-video 2 (related to Figure1 B-I)** The transition from zooids to telotrochs and their subsequent group settlement. Accelerated by a factor of 100. https://youtu.be/ezEXs3lvFgo **Figure 1-video 3 (related to Figure1 B-I)** The transition from zooids to telotrochs and their subsequent group settlement. Accelerated by a factor of 100. https://youtu.be/eQwjoO6vlIM **Figure 1-video 4 (related to Figure1 B-I)** The transition from zooids to telotrochs and their subsequent group settlement. Select segments are accelerated 100-fold. https://youtu.be/EJKMewtThXo **Figure 1-video 5 (related to Figure1 B-I)** The transition from zooids to telotrochs and their subsequent group settlement. Select segments are accelerated 100-fold. https://youtu.be/LKlsDLsH1g0 **Figure 1-video 6 (related to Figure1 B-I)** The transition from zooids to telotrochs and their subsequent group settlement. Select segments are accelerated 100-fold. https://youtu.be/bAN8v2Xz-WM **Figure 1-video 7 (related to Figure1 B-I)** The transition from zooids to telotrochs and their subsequent group settlement without adherence to the substrate. Select segments are accelerated 100-fold. https://youtu.be/zO9UL59LDG8 **Figure 1-video 8 (related to Figure1 J-P)** The transition into telotrochs via binary fission. Select segments are accelerated 100-fold. https://youtu.be/az9qcpWltYk **Figure 1-video 9 (related to Figure1 Q-S)** After swarming, the telotroch settles on a solid surface, secretes a stalk, and reverts into a zooid. Select segments are accelerated 8-fold. https://youtu.be/-C86QUN5fyA **Figure 1-video 10 (related to Figure1)** The majority of the zooids undergo transit without cell division. Select segments are accelerated 100-fold. https://youtu.be/hb9d2x7elTQ **Figure 1-video 11 (related to Figure1 T)** Some individuals transform through cell division while others proceed without it. Select segments are accelerated 100- or 8-fold. https://youtu.be/fD6hb5axIn0 **Figure 1-video 12 (related to Figure1 U)** A few individuals appear to select the settlement location first, with the remaining ones following suit and positioning themselves adjacent to one another. Select segments are accelerated 20.03-fold. https://youtu.be/JetWGHHWKYc **Figure 1-video 13 (related to Figure1 U)** An example of the settlement process. Select segments are accelerated 20.1-fold. https://youtu.be/8dOEmC7t2Zk **Figure 1-video 14 (related to Figure1 U)** An example of the settlement process. Select segments are accelerated 100.06-fold. https://youtu.be/XDPElb5KTqw **Figure 1-video 15 (related to Figure1U)** A representative instance of the settlement process involving a substantial number of telotrochs. Select segments are accelerated 100-fold. https://youtu.be/z-5ND7Cm4JA **Figure 1-video 16 (related to Figure1 V-X)** Two focal points of settlement are established in rapid succession, and the remaining individuals arrange themselves between the two foci as if attempting to avoid gaps. Select segments are accelerated 10-fold. https://youtu.be/1EWbjksuKsQ **Figure 1-video 17 (related to Figure1)** Individuals that arrive later still attempt to settle near the earlier ones, even if those have already begun transforming into zooids. The inset presents an expanded rendition of the upper-right corner. Select segments are accelerated 100- or 7.86-fold. REC/SRC TC: 00:00:00:00/00:00:00:00 indicates the time line time code and source time code in the format hours:minutes:seconds:frames, here and hereafter, unless otherwise explicitly noted. https://youtu.be/knojt56FYc4

We collected submerged fallen leaves from the pond’s bottom and examined them under a compound microscope to locate stalked *V. sp11* (Sanshirou) aggregating densely on the leaves. Leaf fragments were then transferred to an aquarium with aged water containing aquatic plants, such as *Cabomba* sp., or artificial plastic aquatic plants to which the organisms could attach (Figure 1A). It is noteworthy that while they were densely clustered, each organism remained independent. This aggregation on plant fragments was video-recorded at low magnification, and a series of snapshots is provided (Figure 1B-I and figure 1-video 1). The entire population of zooids transitioned to telotrochs almost simultaneously, detaching from their stalks (Figure 1C) and forming a spherical cloud resembling a miniature bee swarm (Figure 1D), as documented by Fox and Newth (1936) for *Vorticella chlorostigma*. The swarm moved about one and a half laps along the inner wall of the well before settling into a dense cluster (Figure 1D-I). Additional examples are presented (Figure 1-videos 2-6). The ITS1-5.8S-ITS2 DNA sequence of *V. chlorostigma* exhibits significant divergence from that of V. sp11 (Sanshirou)(Figure 1-figure supplement 1).

It is noteworthy that the telotroch excretes a gelatinous material, suggesting that the globular cloud is sustained by the mucus as described previously (Fox and Newth, 1936; Horikami and Ishii, 1981). Fox and Newth further identified that a mucous filament originating at the center of the aboral ciliary wreath (described below), which became visible with the addition of Indian ink. This discovery is crucial in deepening our comprehension of the mechanisms driving swarming and/or group settlement behaviors. While we were not aware of this filament during the experiments, we observed turbid material being dragged by the moving globular cloud (arrow in Figure 1F), which became particularly conspicuous under dark-field illumination, potentially arising from the subsequent solidification of the mucus fibers. At times, this material was incorporated into the settled cluster (arrow in Figure 1H). They additionally observed: ‘At the end of swarming they are so solid that sometimes groups of vorticellids settle down on them as they would on pondweed.’ (Fox and Newth, 1936). A video illustrating the formation of telotroch clusters settling without adherence to the substrate (Figure 1-video 7) may exemplify the phenomenon described. The formation of the mucus fibers will be addressed in more detail later.

### Life cycle of Vorticella

The transition into telotrochs via binary fission was video-recorded at higher magnification (Figure 1J-P and Figure 1-video 8). The zooid, equipped with a prominent wreath of cilia at its oral end (oral cilia) used for feeding (Figure 1J), becomes more rounded (Figure 1K), initiates division (Figure 1L), and forms two daughter cells (Figure 1M). The oral cilia retract, while the aboral end becomes encircled by a wreath of cilia (aboral cilia) that emerge from aborally positioned kinetosomes (Buhse, 1998) (Figure 1M). The cell elongates, transforms into a telotroch, detaches from the stalk, and swims with the aboral end leading, propelled by the aboral cilia (Figure 1M-P). After swarming, the telotroch settles on a solid surface, secretes a stalk, and reverts into a zooid (Figure 1Q-S and Figure 1-video 9). Not all cells undergo division prior to transformation. In certain samples, the majority undergo transit without cell division (Figure 1-video 10), whereas others exhibit a combination of both types [Figure 1-video 11 and Figure 1T, some individuals transform through cell division (..) while others proceed without it (.)]. The genotypes of both populations are identical. As described later, this transformation appears to occur daily, with cell division taking place on the first day but not on the following day in the observed case, indicating that the presence or absence of division is driven by physiological conditions, such as nutrition, rather than genetic inheritance. Despite the presence of both modes within the population, the timing of the transformation post-division is not significantly delayed compared to that of the population without division (Figure 1-video 11), suggesting a highly regulated transformation mechanism across individuals.

It is interesting to explore how and when telotrochs decide on a site for settlement. A few individuals appear to select the settlement location first, with the remaining ones following suit and positioning themselves adjacent to one another (Figure 1U and Figure 1-video 12, additional examples Figure 1-video 13-15). The decision-making process is not straightforward; they continue roaming, searching for a site, and may even communicate with each other via physical contact. In another instance, when numerous telotrochs were involved, two focal points of settlement were established in rapid succession, and the remaining individuals arranged themselves between the two foci as if attempting to avoid gaps (Figure 1V-X and Figure 1-video 16). An additional example underscores another facet of the behavior: later-arriving individuals persist in seeking proximity to those that preceded them, even when a substantial number have already commenced their transformation into zooids (Figure 1-video 17). This suggests a strong preference for settling in as densely packed a manner as possible.

We conducted prolonged observations exceeding 60 hours and determined that the transformation cycle from zooid to telotroch occurs at intervals of approximately 21 hours. The telotrochs congregated at location 1 (Figure 2 and Figure 2-video 1) and swarmed toward location 2, and so forth. The intervals between the initiation of each swarming event were approximately 20 hours (from swarming away from 1 to swarming away from 2), 20.5 hours (from 2 to 3), and 23 hours (from 3 to 4). Hence, the average interval between successive swarms was roughly 21 hours. This observation aligns with the previous report indicating an interval of around 24 hours (Fox and Newth, 1936).

**Figure 2.**
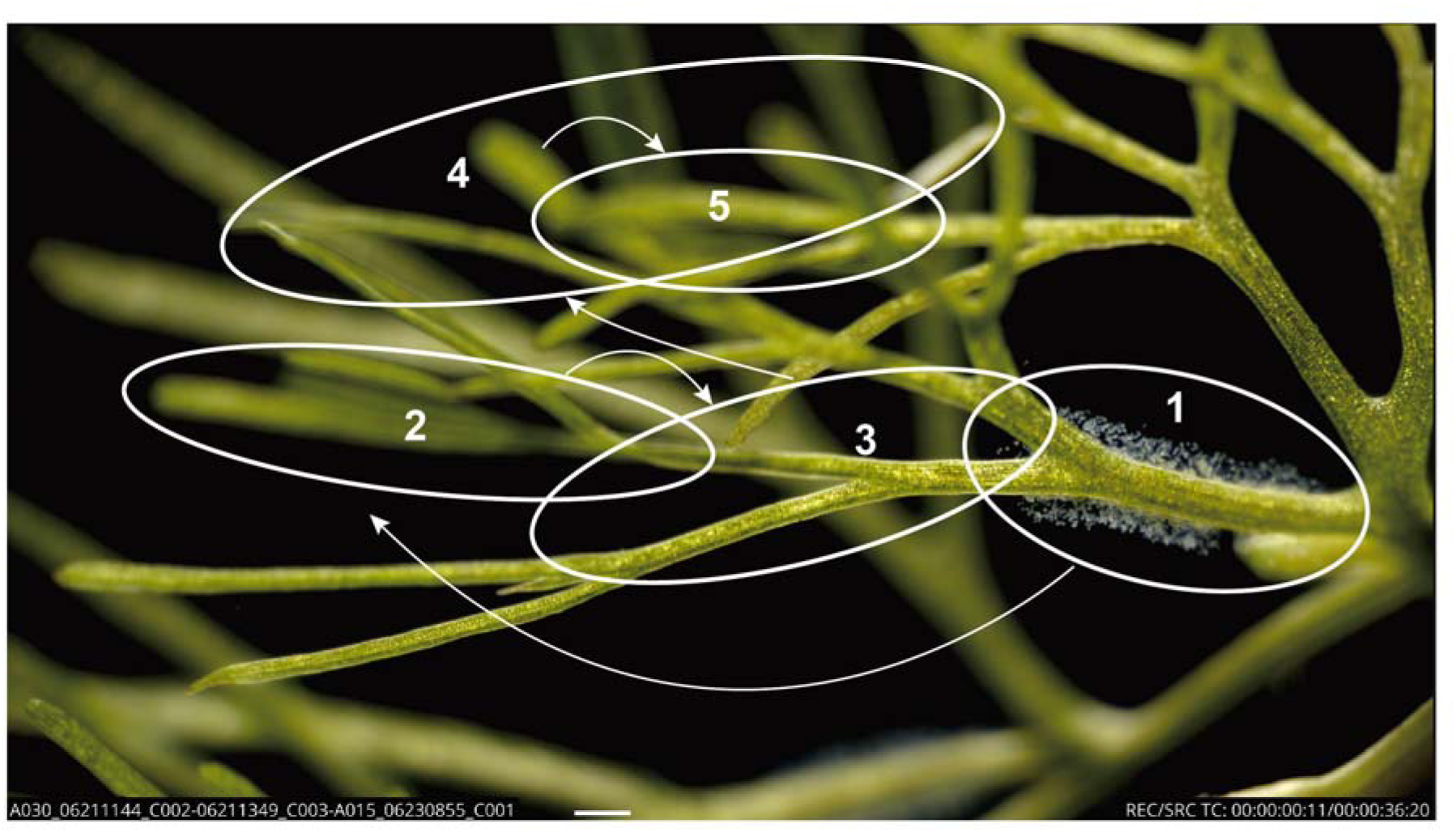
The swarming behaviors documented through video recordings spanning 60 hours. The settlement location transitioned sequentially from positions 1 to 5, with each shift occurring at an interval of approximately 21 hours. Scale indicates 1 mm. **Figure 2-video 1 (related to Figure2)** Behaviors were recorded at 30 frames per second for the initial 2 hours and at 1 frame per second for the remaining duration. The recording length is approximate, as SSD storage swaps were conducted for extended recording periods. All source files were concatenated into a single compound file for editing purposes. The source time code format 01:01:01:01 represents 30 hours:0.5 hours:0.5 minutes:1 second, with each 30-second increment advancing the format by one unit. Select segments are accelerated 120-, 240- or 3000-fold. https://youtu.be/1BrYJ43Uo88

### Mucous fiber formation

The existence of mucus is discernible upon the mechanical displacement of the swarm (Figure 3A-B and Figure 3-video 1), as articulated by Fox and Newth, who noted that ’the presence of mucus would not be suspected but for the fact that the mass, which has a gelatinous consistency, can be pulled aside bodily with a needle’(Fox and Newth, 1936) Under dark-field illumination (Figure 3C-K), an indistinct thread—presumably composed of mucous fibers, as inferred from their findings—emerges shortly after the swarm commences (Figure 3E). The thread becomes increasingly defined, thickens, and ultimately condenses into a globular structure (Figure 3F-J), consistent with their observation that ‘the movements of the animals themselves may cause the progressive solidification of the mucus which occurs during swarming.’ (Fox and Newth, 1936). It follows the swarm until settlement on the substrate but does not necessarily participate in the settled population, as observed in this particular instance. The condensed mucous fiber was attached above the plant fragment, while the telotrochs settled beneath, out of focus (Figure 3K). We lacked the opportunity to stain the mucous fiber for closer examination and thus cannot describe its characteristics in detail, leaving it unclear whether the telotrochs retain the mucous fibers post-condensation at the end of swarming.

**Figure 3.**
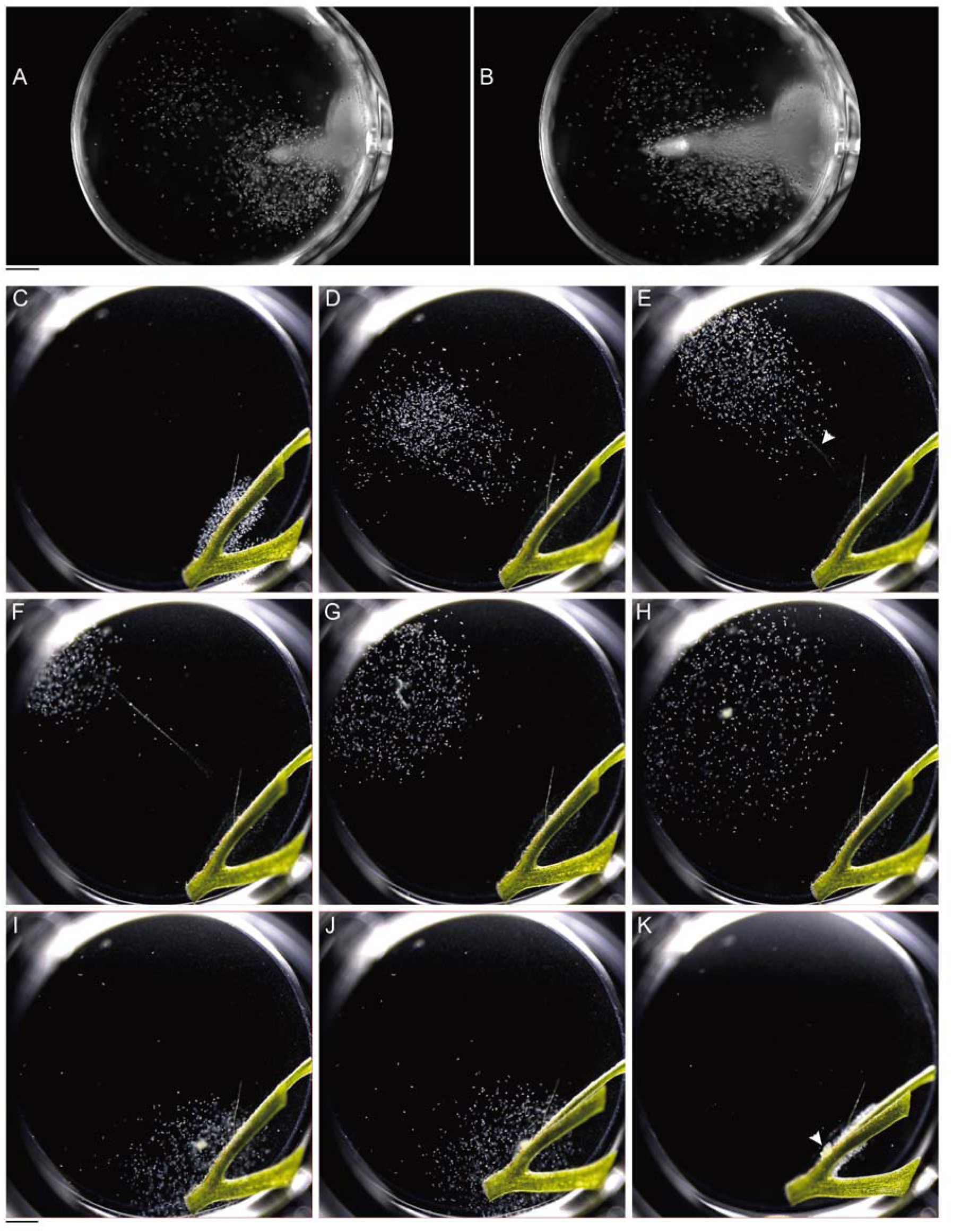
Telotrochs secrete mucus. (A-B) A spherical telotroch swarm being drawn aside bodily with a needle. (C-K) The progression of mucous fiber development accompanying telotroch swarming. An arrow denotes the faint, early-stage mucous fiber (E), while an arrow indicates the condensed mucous fiber positioned apart from the settled telotrochs (K). Scales indicate 1 mm. **Figure 3-video 1 (related to Figure3A-B)** Select segments are accelerated 100.06-fold. https://youtu.be/TTx5ickofxs **Figure 3-video 2 (related to Figure3C-K)** Accelerated by a factor of 100. https://youtu.be/iLhBgE3D0kk

### What drives their propensity to swarm and congregate in groups upon settlement?

What is benefit of this behavior? In unicellular organisms, this phenomenon can be interpreted from an evolutionary standpoint as the origin of multicellularity. A notable example is the slime mold Dictyostelium discoideum, which proliferates as free-living unicellular amoebae through cell division and, upon starvation, aggregates to construct a multicellular fruiting body with differentiated cysts or spores. Organisms displaying such life cycles have independently evolved at least six times across both eukaryotic and prokaryotic lineages, including fungi and ciliates (Du et al., 2015). However, in the case of *Vorticella*, they remain independent, exhibiting swarming and spatial proximity without coalescing into a multicellular structure akin to a “slime mold.” One hypothesis is that their close assemblage may facilitate sexual conjugation, though we have yet to observe such events as previously noted (Fox and Newth, 1936). Recent investigations into *Stentor coeruleus*, a suspension-feeding unicellular ciliate like *Vorticella*, which exists both solitarily and colonially, suggest that hydrodynamic interactions between neighboring individuals enhance feeding efficiency through accelerated flow, contingent on inter-individual spacing (Shekhar et al., 2023). This could potentially account for the pronounced clustering observed in these ciliates. However, if this hypothesis is valid, then why does *Vorticella* repeatedly undergo cycles of aggregation and dispersion daily, despite the significant energy expenditure required for the transition between zooid and telotroch forms? A potential explanation for this behavior may lie in their predator evasion strategy, as outlined below.

We sporadically observed the predatory ciliate, *Amphileptus sp*., feeding on *Vorticella* (Figure 4 A-D and Figure 4-video 1). Individuals within a group can be consumed sequentially if they remain in their stalked sessile form (Figure 4 E-I and Figure4-video 2). The periodic transition to the telotroch stage and relocation may serve as a strategy to evade predators, while swarming enables them to regroup for more efficient feeding.

**Figure 4.**
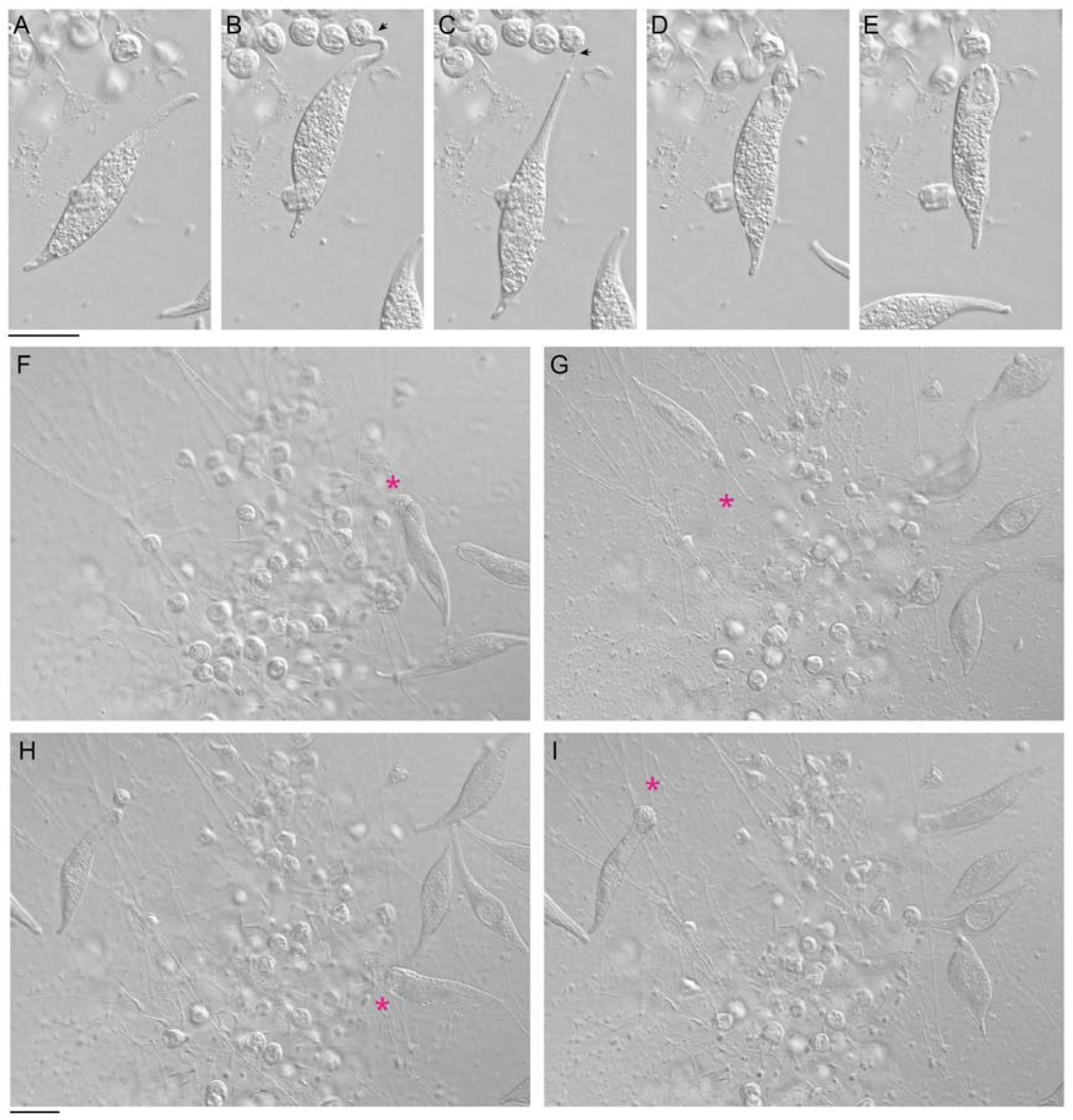
A predatory ciliate, *Amphileptus sp*., preying upon Vorticella. (A-E) The intricate process by which *Amphileptus sp.* preys upon *Vorticella sp11 (Sanshirou)* was observed. Upon mechanical contact with *Vorticella* using its proboscis (B, arrow), *Amphileptus* appears to discharge prexicysts or analogous organelles (C, arrow), akin to the behavior exhibited by Didinium (Buonanno and Ortenzi, 2021) facilitating the capture of *Vorticella*. (F-I) A cluster of *Vorticella sp11 (Sanshirou)* was sequentially consumed by a group of *Amphileptus sp* (an asterisk). **Figure 4-video 1 (related to Figure 4A-E)** Select segments are accelerated 8-fold. https://youtu.be/nzoriPLGvcI **Figure 4-video 2 (related to Figure 4A-D)** The instance of Amphileptus sp. preying upon Vorticella is indicated by an asterisk. Select segments are accelerated 100-, 20.1-, 19.97- or 8-fold. https://youtu.be/vZD5u9RqMDg

### The case of *V. campanula*

We also identified a distinct species of *Vorticella*, tentatively designated as *V. cf. campanula* (Sanshirou), due to its morphological resemblance to *V. campanula* (notably, the shape of the cell body and the presence of refractive oil droplets within the cytoplasm, which give the cells a darkened appearance) (Figure 4), as well as its genomic sequence. The ITS1-5.8S-ITS2 region of *V. cf. campanula* (Sanshirou) exhibits the highest sequence homology to that of *V. campanula*, despite some degree of heterogeneity observed among individual specimens (Figure 4-supplementary figure1). The behaviors of *V. cf. campanula* (Sanshirou) have not been extensively studied in comparison to *V. sp11* (Sanshirou), yet they exhibit significant similarities with previous research on *V. campanula* (Fox and Newth, 1936; Fox and NEWTH, 1935). *V. cf. campanula* (Sanshirou) transitions from the zooid to telotroch stage (Figure 5A-D) and settle in groups (Figure 5E-G), much like *V. sp11* (Sanshirou). The most pronounced difference lies in their swarming behavior. *V. sp11* (Sanshirou) forms a compact, ball-like cluster resembling a cloud of bees, whereas *V. cf. campanula* (Sanshirou) moves back and forth along mucous threads (Figure 5H), as shown by the line from the upper left corner to the middle bottom, along which individuals slide (Figure 5H). In more natural conditions, these threads can extend up to 10 cm in length (Fox and Newth, 1936; Fox and NEWTH, 1935). The swarming behaviors of the two species were compared (Figure 5I-J). *Vorticella sp11* (Sanshirou) exhibited a shorter swarming duration, forming a compact and dense cloud and ultimately settling into a tightly clustered group (Figure 5I) as described. In contrast, *Vorticella cf. campanula* (Sanshirou) moved along a trajectory for an extended period before settling in smaller, more dispersed groups (Figure 5J). When *V. cf. campanula* (Sanshirou) transitioned to telotrochs in a larger container, swarming units initially appeared locally, gradually merging and morphing over time, and eventually coalescing into a unified, larger stream (Figure 5K-N). This example illustrates the swarming behavior across a more extensive area, paralleling previous observations (Fox and Newth, 1936; Fox and NEWTH, 1935), despite the inherent differences between our two-dimensional experimental conditions and their more natural, three-dimensional environment, which was described as follows: ’Numerous individuals swim close to the mucous thread-some moving round it, some swimming upwards, others downwards-while all the time individuals leave the thread and return to it after short horizontal excursions.’(Fox and Newth, 1936)

**Figure 5.**
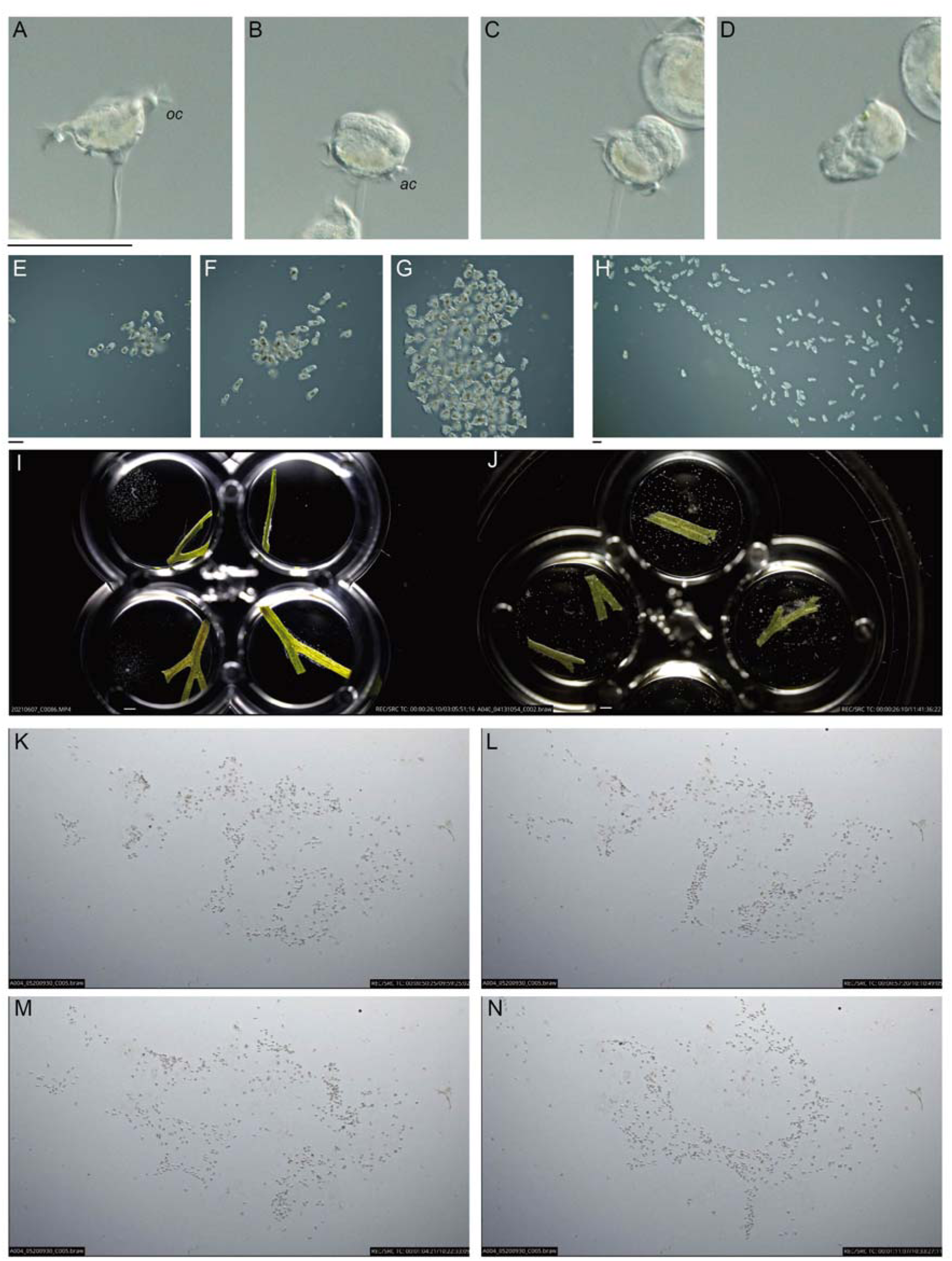
Swarming behaviors of *V. cf. campanula (Sanshirou).* (A-D) Transition from zooids to telotrochs. (A) *oc* denotes oral cilia, while (B) *ac* indicates aboral cilia. (E-G) Settlement of telotrochs. (H) Telotroch swarming. (I-J) Side-by-side comparison of swarming behaviors between *V. sp11 (Sanshirou)* (The upper-left specimen is previously exhibited in Figure 3C-K) (I) and *V. cf. campanula (Sanshirou)* (J). Scales indicate 1 mm. (K-N) Telotroch swarming coalescing and forming a single, unified stream. Scales are not available. **Figure 5-video 1 (related to Figure 5A-D)** Select segments are accelerated 50.18- or 50.01-fold. https://youtu.be/DKpSIFoM1YU **Figure 5-video 2 (related to Figure 5E-G)** Select segments are accelerated 20.09-fold. https://youtu.be/9zkQYLnVqk8 **Figure 5-video 3 (related to Figure 5H)** https://youtu.be/9gwk3RgWn4M **Figure 5-video 4 (related to Figure 5I-J)** The two video recordings (I and J) were conducted independently and integrated during the editing process. Accelerated 20.09-fold. https://youtu.be/0wlM_yDZvzY **Figure 5-video 5 (related to Figure 5K-N)** The recording field expanded progressively as the swarm’s area increased. Select segments are accelerated 100.11-fold. https://youtu.be/ZkqdtnE0GDA

The analogous swarming behaviors of the two types of *Vorticella* have been documented, nearly a century apart, in Great Britain (Fox and Newth, 1936; Fox and NEWTH, 1935)and Japan (this study). Certain *Vorticella* species do not aggregate but instead settle solitarily on surfaces. Horikami and Ishii, in their conference abstract, reported that *V. campanula*, *V. convallaria*, *V. monilata*, and *V. vestita* exhibited group settlement behaviors.

Additionally, they described that a viscous substance could attract telotrochs in a species-specific manner (Horikami and Ishii, 1981). The role of mucus in swarm formation remains uncertain. Does it merely encapsulate the telotrochs to restrict their mobility, or does it possess a physiological function that modulates telotroch behavior, as indicated by Horikami and Ishii? Is the strand observed by Fox and Newth derived from the mucus, contributing to the formation of the putative condensed mucus-like material we detected? Regrettably, we lack evidence to address this question, as we only discovered the relevant publication on mucus fibers after completing our experiments. We are also intrigued by the duration for which the mucus exerts its influence during the settlement process. In certain instances (e.g., Figure 3-video 2), the condensed mucus-like material does not participate in the settlement cluster. Conversely, upon closer examination of the settlement process (e.g., Figure 1-video 12), a few telotrochs initially rotate around the condensed mucus-like material, appearing to be constrained by a central tension, suggesting a persistent physical linkage. Elucidating the molecular function of this mucus, along with any potential swarming behaviors exhibited by other *Vorticella* species, would present an intriguing direction for future research.

If the dense clustering observed in this *Vorticella* settlement is not considered the precursor to multicellularity, then it may represent an evolutionary origin of coordinated group actions in multicellular organisms. Eduardo Orias posited that ciliates could be perceived as multicellular organisms. He proposed that ciliates with multiple cilia evolved from flagellates bearing a single flagellum through a mutation that delayed cytokinesis, resulting in chains of flagellates(Orias, 1976) . In this context, ciliates can already be viewed as multicellular organisms, and the *Vorticella* species described here could be characterized as “gregarious species”, as previously noted (Fox and Newth, 1936).

## Materials and Methods

### Identification of the *Vorticella* species

Genomic DNA was isolated from 5-10 zooids and the ITS1–5.8S–ITS2 region was amplified by polymerase chain reaction (PCR) using the SB2 (5′-GTAGGTGAACCTGCGGAAGGATCATTA-3′) (Goggin and Murphy, 2000)and ITS-R (5′-TACTGATATGCTTAAGTTCAGCGG-3′) (Sun et al., 2010). The thermal cycling conditions were as follows: initial denaturation at 94°C for 2 minutes, followed by 35 cycles of denaturation at 98°C for 10 seconds, annealing at 60°C for 30 seconds, and extension at 72°C for 60 seconds, with a final extension step at 72°C for 7 minutes. The DNA database was queried for sequence homology utilizing the Basic Local Alignment Search Tool (BLAST) available at https://blast.ncbi.nlm.nih.gov/Blast.cgi. The DNA sequences were aligned utilizing the ApE Plasmid Editor v3.1.4 (Wayne Davis) and edited using Illustrator (Adobe).

### Microscopy and video recordings

The organisms were examined using the Axio Observer, Axiovert, and Axio Zoom V.16 microscopes (Carl Zeiss). Video recordings were captured with the URSA Mini Pro 12K [Blackmagic Design (BMD)], Blackmagic Pocket Cinema Camera 6K (BMD), ILCE-1 (Sony), and S1H (Panasonic) cameras, and subsequently edited using DaVinci Resolve 19 (BMD).

The manuscript underwent partial grammatical refinement using ChatGPT.

## Acknowledgements

We thank Toshinobu Suzaki for providing a conference abstract and Ryu Ueda for helpful comments. We are grateful to Minoru Saitoe and Takashi-Adachi Yamada for their generous supports. This work was supported by grants from the Ministry of Education, Culture, Sports, Science and Technology (21K19288) to T.T.

## Author contributions

T.T. designed the research, performed imaging and wrote the manuscript. Y.M. performed the PCR analysis.

## Competing interest

The authors declare that no competing interests exist.

Additional files

Supplementary files

Figure 1-supplementary figure1

Figure 4-supplementary figure1

