## Supplementary material for "The swarming behaviors of Vorticella": Figure 1-supplementary figure1 (from another version of this preprint: version 1, 2024-11-28)

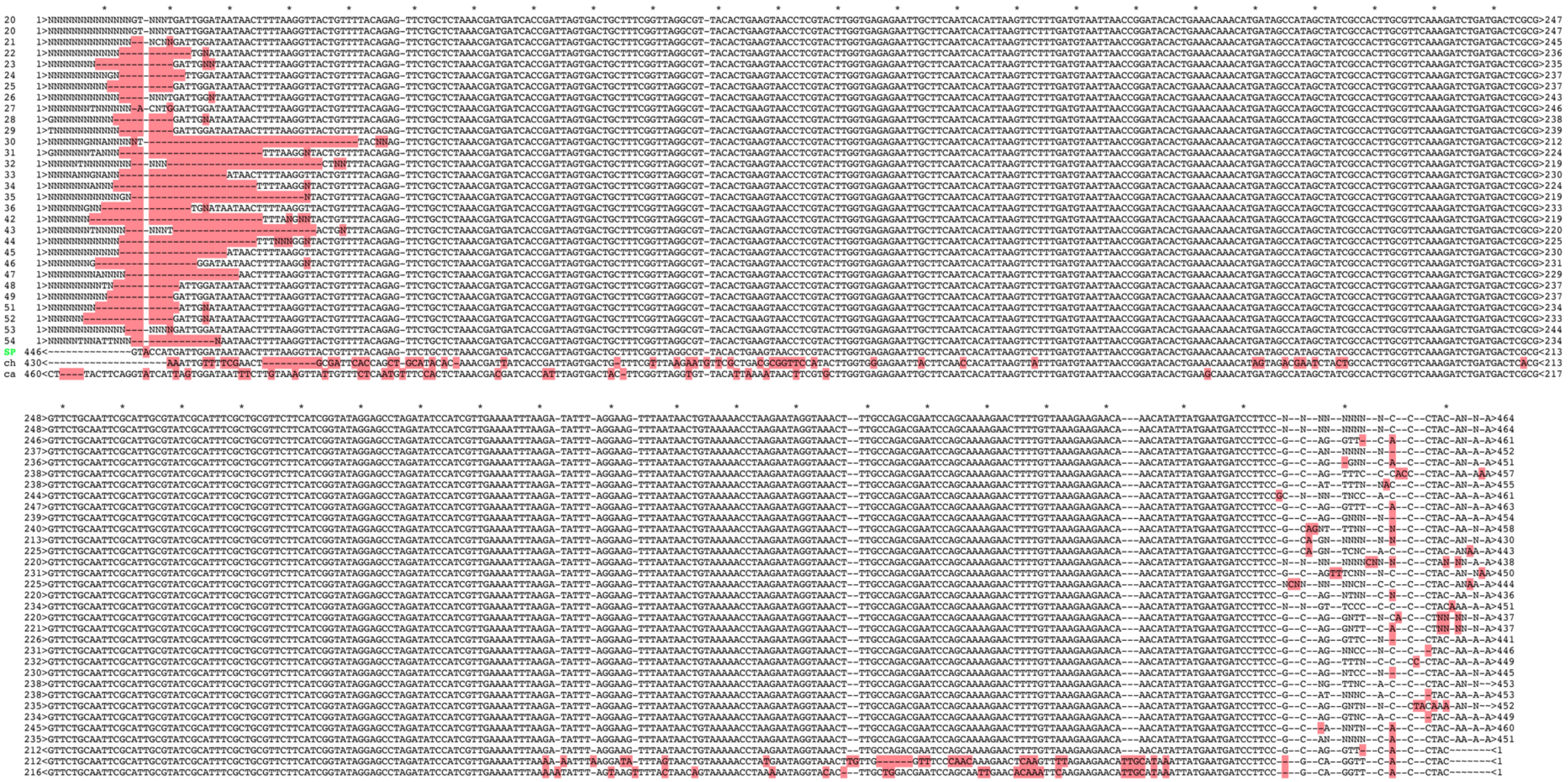

The ITS1-5.8S-ITS2 DNA sequence of the initial group of *Vorticella* obtained from Sanshirou-ike (sample number 20) was aligned with sequences from *Vorticella* specimens collected from Sanshirou-ike (sample numbers 20–54), *V. sp11* (SP), *V. chlorostigma*(ch), and *V. campanula* (ca). The DNA sequences distinct from that of specimen number 20 are highlighted in red. Sample numbers corresponding to video-recorded specimens are indicated in parentheses: Figure 1-video 1 (26), Figure 1-video 4 (45), Figure 1-video 5 (48), Figure 1-video7 (46), Figure 1-video8 (27), Figure 1-video10 (20), Figure 1-video11 (29), Figure 1-video15 (18) and Figure 1-video17 (21).
