## Supplementary material for "The swarming behaviors of Vorticella": Figure 4-supplementary figure1 (from another version of this preprint: version 1, 2024-11-28)

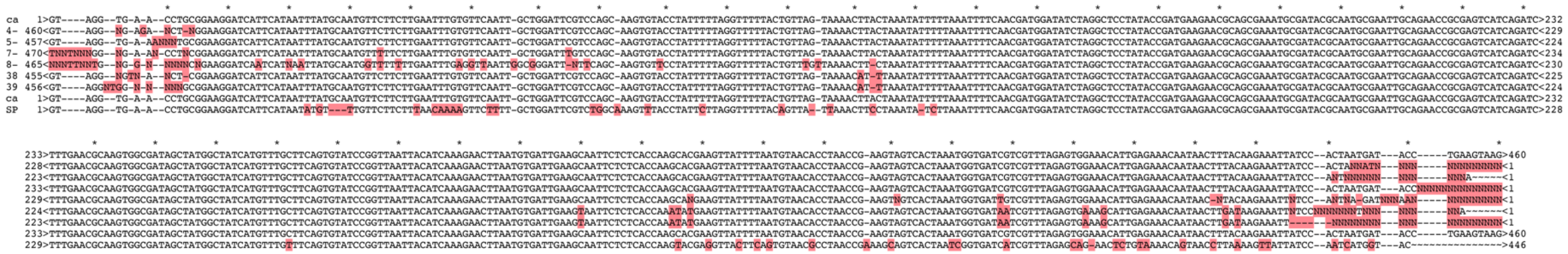

The ITS1-5.8S-ITS2 DNA sequence of *V. campanula* (ca) is aligned with sequences from the second group of *Vorticella* specimens collected from Sanshirou-ike (sample numbers 4–39), *V. campanula* (ca), and *V. sp11* (SP). The DNA sequenceces distinct from that of *V. campanula* are highlighted in a reddish hue. The DNA sequence of the specimens, whose video recordings are presented in Figure 5, corresponds to No. 7 or 8.
